# Establishment of murine *in vitro* blood-brain barrier models using immortalized cell lines: co-cultures of brain endothelial cells, astrocytes, and neurons

**DOI:** 10.1101/435990

**Authors:** Fakhriedzwan Idris, Siti Hanna Muharram, Zainun Zaini, Suwarni Diah

## Abstract

Blood-brain barrier (BBB) is a selective barrier formed by the endothelial cells that line cerebral microvessels. It serves as a physical barrier due to the presence of complex tight junctions between adjacent endothelial cells which limits the paracellular movement of most molecules across the BBB. Many *in vitro* models of the BBB have been established to mimic these *in vivo* properties with limited success. In this study, we described the properties of a cell-based murine *in vitro* BBB model in five configurations constructed using immortalized cell lines in a 12-well format Transwell system: murine brain endothelial cells (bEnd.3) grown in a monoculture, or as co-culture in contact with astrocytes, or without contact with astrocytes or neurons, and triple co-culture combining the three cell lines. We found that only contact and triple co-culture model closely mimic the *in vivo* BBB tightness as evaluated by apparent permeability (Papp) of sucrose and albumin producing the lowest Papp values of 0.56 ± 0.16 × 10^−6^ cms^−1^ and 3.30 ± 0.51 × 10^−6^ cms^−1^, respectively, obtained in triple co-culture model. Co-culturing of bEnd.3 with astrocytes increased the expression of occludin as shown by western blot analysis, and immunohistochemistry showed an increase in peripheral localization of occludin and claudin-5. In addition, we found conditioned media were able to increase *in vitro* BBB model tightness through the modulation of tight junction proteins localization. We conclude that the presence of astrocytes and neurons in close proximity to brain endothelial cells is essential to produce a tight *in vitro* BBB model.

## 1. Introduction

The blood-brain barrier (BBB) is a selective barrier and an intricate structure comprising of microvascular endothelial cells (ECs) that made up the vessel wall, in close association with basement membrane or basal lamina and other cells including pericytes, astrocyte end-feet, microglia and neurons to form an organization known as the neurovascular unit (NVU) (Risau and Wolburg, 1990; Begley and Brightman, 2003). It serves as a physical barrier due to the presence of complex tight junctions (TJ) between adjacent endothelial cells (Begley and Brightman, 2003; Watanabe et al., 2013) which force most molecular traffic to take a transcellular route across the BBB, instead of moving paracellularly through the junctions, as in most endothelia (Wolburg and Lippoldt, 2002; Hawkins and Davis, 2005).

*In vitro* models of the BBB were developed in order to study further the structure, physiology, and pathology of the BBB. These models utilized primary brain capillary endothelial cells extracted from porcine, bovine, rat, mouse and human tissues which demonstrated a wide range of properties (Perrière et al., 2005; Malina et al., 2009; Nakhlband and Omidi, 2011; Vandenhaute et al., 2011; Cantrill et al., 2012; Patabendige et al., 2013). Cell lines have also been used in the establishment of *in vitro* BBB models in addition to primary cells, such as HBMEC or hCMEC/D3 which express markers of brain endothelial cells but show low transendothelial electrical resistance (TEER) (Weksler et al., 2005; Förster et al., 2008; Hatherell et al., 2011; Eigenmann et al., 2013). Stem cells have been used recently to produce human brain ECs including human pluripotent stem cells (hPSCc) (Lippmann et al., 2012) and human cord blood-derived stem cells of circulating endothelial progenitor and hematopoietic lineages (Boyer-Di Ponio et al., 2014; Cecchelli et al., 2014). These stem cells-derived ECs developed restrictive barrier with well-formed TJ proteins. Most models are developed as a monolayer which is grown on a microporous membrane filter culture insert. However, astrocytes have been found to play important roles in the establishment and development of BBB phenotypes (Arthur et al., 1987; Janzer and Raff, 1987) which led to the development of co-culture model combining endothelial cells and astrocytes or endothelial cells grown in astrocyte-conditioned media (Terasaki et al., 2003). In this study, we utilized immortalized cell lines instead of primary cells as cell lines possess several advantages; 1) reliability, in which cells can be obtained and purchased from established sources; 2) consistency as the variability can be reduced or eliminated between different sets of experiments since the source of cells is controlled; 3) longevity and durability since cell lines are immortalized by a transfection process which may preserve the important cell features and prevent the loss of these features over time (Levashova et al., 2003); 4) efficiency in terms of the preparation time and cost which are reduced as cells can be used immediately upon purchase. Despite these advantages, BBB models utilizing immortalized cell lines are generally leakier compared to primary cultures, which restricts their use for permeability screenings. Although most cell lines preserve basic cerebral endothelium-like features, the immortalization process may affect the expression level of enzymes, TJs, and transporters from the SLC and ABC families (Urich et al., 2012). Nevertheless, cell lines have proved useful for other studies, for example biochemical and mechanistic studies of the BBB and for drug delivery systems targeting the central nervous system (Reichel et al., 2003; Roux and Couraud, 2005).

Developing murine *in vitro* BBB models is necessary as these models are potentially better surrogates to correlate with *in vivo* data since many *in vivo* studies especially on drug transport across BBB are done in mouse models. Furthermore, using murine BBB models may be more suitable in BBB studies involving neurodegenerative disorders, brain cancer, and inflammatory event because these diseases are commonly investigated in mouse models (Shayan et al., 2011). Our study was aimed to establish and compare five murine *in vitro* BBB models constructed using immortalized cell lines grown on Transwell inserts: a monoculture, a contact co-culture with astrocytes, two non-contact co-cultures, and a triple co-culture. We also determined the effects of conditioned media obtained from astrocytes and neurons on bEnd.3 cells.

## 2. Results

### 2.1. bEnd.3 (mouse Balb/c), astrocyte type III (C8-D30) (mouse C57BL/6), and neuroblastoma (SH-SY5Y) (human) identification and characterization *in vitro*

bEnd.3, C8-D30, and SH-SY5Y cells proliferated well on tissue culture flask under routine maintenance. Figure 1 shows bEnd.3 cells at passage 2, C8-D30 and SH-SY5Y at passage 8.

**Figure 1.**
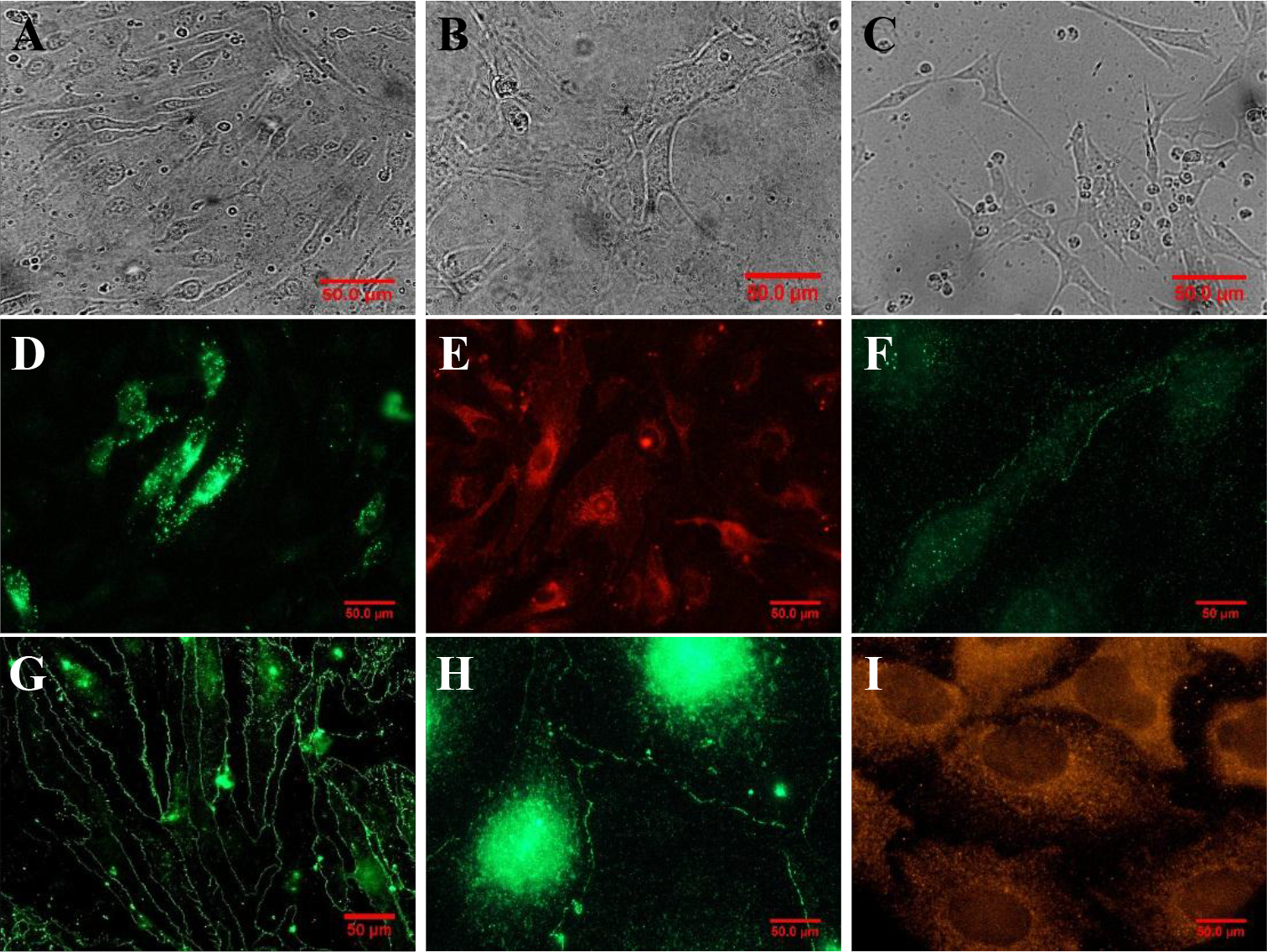
Identification and characterization of brain cells. Different cells showed unique morphologies which can be used as identification; A) bEnd.3 with spindle-shaped, B) C8-D30 with star-shaped, and C) SH-SY5Y with neuroblast-like morphology. bEnd.3 was further characterized by the presence of D) von Willebrand factor and positively stained with (E) LEA. The detection of TJ proteins including F) occludin, G) claudin-5, and H) ZO-1 at the cell-cell contacts further confirmed bEnd.3 as brain microvascular endothelial cells. C8-D30 lineage was confirmed by the detection of I) GFAP.

bEnd.3 was recognizable by the relatively large size of the cells with spindle-like morphology and positively stained for von Willebrand factor, TJ proteins: occludin, claudin-5, and zonula occludens-1 (ZO-1), in addition to the staining with tomato lectin (LEA). While C8-D30 showed a star shape morphology with short astrocytic fibers and positively stained for glial fibrillary acidic protein (GFAP). SH-SY5Y was recognizable by their morphologies which were described as neuroblast-like cell line (N-type) with observable dendrites. von Willebrand factor, occludin, claudin-5, ZO-1, and GFAP were not detected on SH-SY5Y cells. During routine maintenance of cells, SH-SY5Y was observed to grow as a mixture of floating and adherent cells, while, bEnd.3 and C8-D30 grow as adherent cells.

### 2.2. Permeability assessment of established *in vitro* BBB models

To examine the effect of different cell types on bEnd.3 cell integrity, different sets of BBB co-cultures were established and their paracellular and transcellular permeability was assessed using transport study. We investigated the influence of immortalized astrocytes and neurons derived from mouse and human, respectively, on our murine endothelial cell model. As shown in Figure 2, the paracellular and transcellular permeability of bEnd.3 monolayers quantified by sucrose and albumin was the highest (96.1 ± 24.5 and 22.9 ± 7.5 × 10^−6^ cms^−1^, respectively). Compared to bEnd.3 monolayers with no influence of other cells (monoculture), all co-culture models exhibited significantly lower permeability for sucrose and albumin. There was no significant difference in sucrose permeability in the non-contact models containing bEnd.3/C8-D30 cells and bEnd.3/SH-SY5Y cells. The effect was greater in triple co-culture model in which C8-D30 cells were positioned on the basal side of the inserts and SH-SY5Y on the bottom of the well, with a significant 172.2-fold decrease in Papp of sucrose. Nonetheless, the effects of different co-culture models on sucrose permeability were not significant. While, the transcellular permeability measured using albumin revealed only contact and triple co-culture models showed significant reduction as high as 3.8- and 3.7- fold, respectively, compared to the monoculture. A high albumin permeability across the BBB models may be attributed by the increased transport of albumin via endocytosis or transcytosis processes mediated by caveolae present on surfaces of endothelial cells (Li et al., 2013). These processes are temperature-dependent as described by Smith and Borchardt (1989) and Li et al. (2013); higher temperature (37°C) enhanced the transcytosis activity of albumin across endothelial cells.

**Figure 2.**
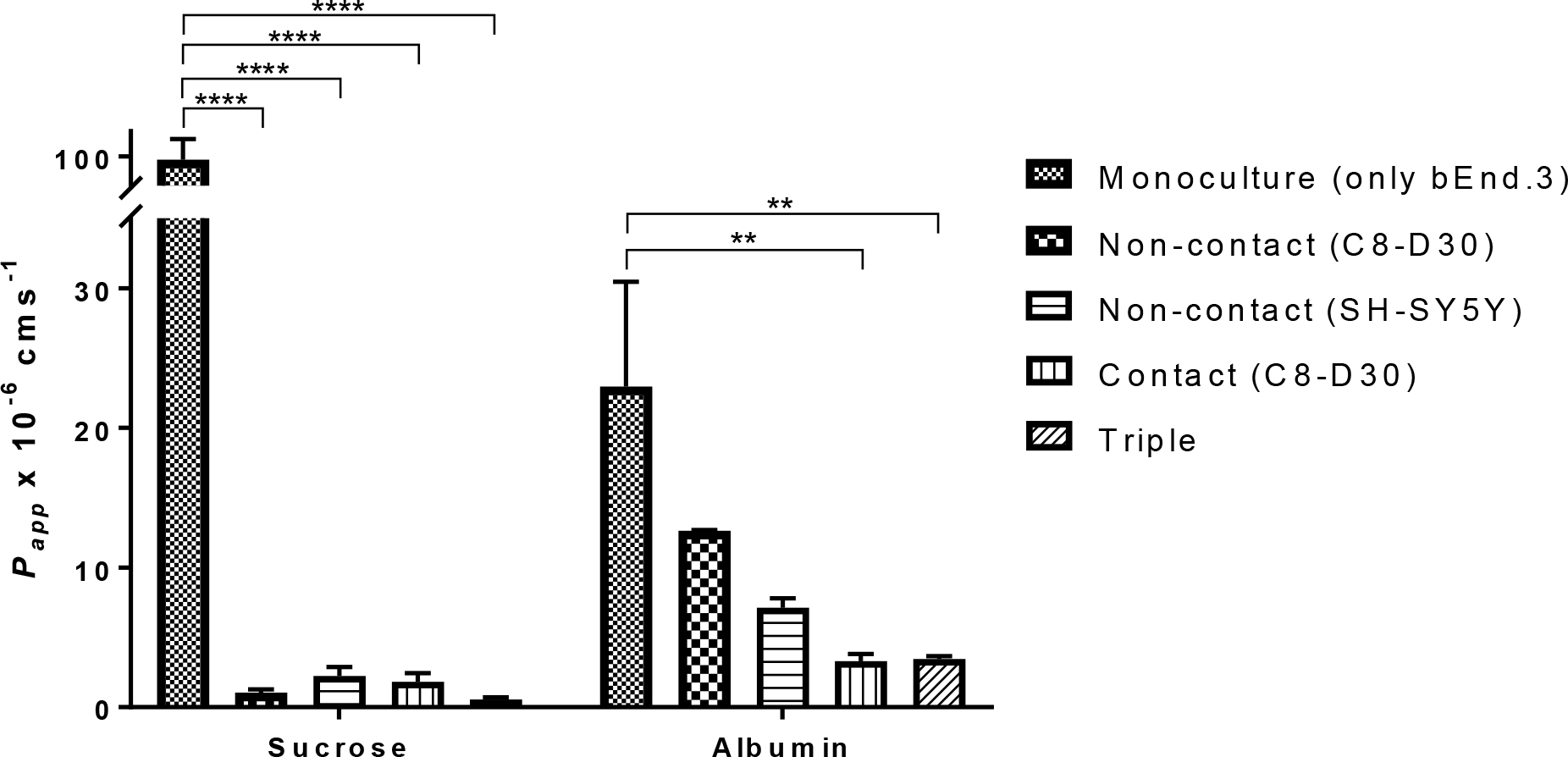
Permeability assessments using transport study on different *in vitro* BBB models. Two standard markers were used: sucrose (paracellular marker) and albumin (transcellular marker). Data are presented as means ± SD (*n* = 20 – 24) from 3 independent experiments. Asterisks indicate significance (** *p*<0.01, **** *p*<0.0001).

### 2.3. Regulation of TJ proteins expression in different *in vitro* BBB model set-ups

Paracellular permeability of BBB is functionally associated to the junctional molecules expression. Hence, to elucidate the mechanism of decreased permeability in different culture models, we performed protein expression analysis by quantifying specific TJ proteins which are known to play roles in the regulation of endothelial TJs. Here, we studied three proteins namely occludin, claudin-5, and ZO-1. We performed Western blotting to quantify the levels of TJ proteins in bEnd.3 cells cultured in the presence or absence of C8-D30 and SH-SY5Y. As shown in Figure 3, the presence of these cells relatively increased the expression of endothelial TJ proteins with the highest seen in contact and triple co-cultures. However, significant increase level was only observed for occludin in the contact co-culture set-up with C8-D30 as high as 4.5- fold, but not the others compared to the monoculture. Co-culturing bEnd.3 with C8-D30 and SH-SY5Y regardless of the set-ups did not significantly elevate the levels of claudin-5 and ZO-1.

**Figure 3.**
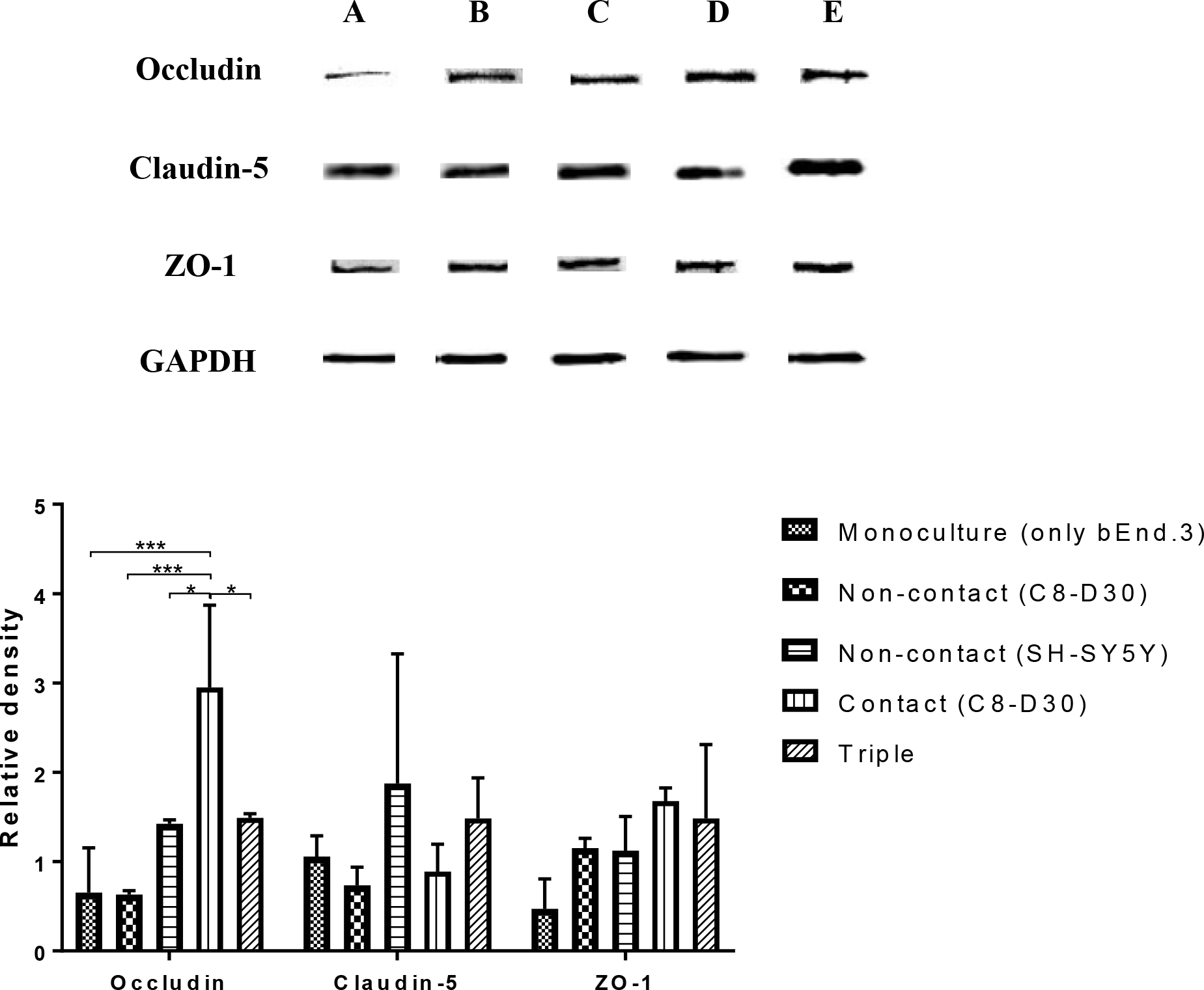
Expression of proteins determined by Western blotting. A) Monoculture (only bEnd.3), B) non-contact (bEnd.3/C8-D30), C) non-contact (bEnd.3/SH-SY5Y, D) contact model (C8-D30 positioned at the basal side of insert), and E) triple co-culture. The relative density with GAPDH as the reference was quantified using ImageJ software. Asterisks indicate significance (* *p*<0.05, *** *p*<0.001).

### 2.4 Effects of co-culturing on the peripheral distribution of TJ proteins in bEnd.3 cells

Microscopic examination of bEnd.3 monolayers by immunohistochemistry showed that all models exhibited staining for junctional proteins occludin, claudin-5, and ZO-1 (Figure 4). Peripheral protein detection was important to determine the distribution of these proteins within the cells which contributes to the tightness of developed models. Upon observation of bEnd.3 grown in a monoculture, only some cells showed peripheral localization of occludin and most were detected in the cytoplasm as shown by the high intensity especially near and around the nuclei. The introduction of C8-D30 or SH-SY5Y into the culture systems increased the peripheral localization of occludin at cell-cell contacts. Further analysis using ImageJ software of the peripheral intensities (Figure 4D) showed a significant increase in all culture models compared to the monoculture. It was especially high for contact model which showed 5.9-fold increase in intensity and significantly different from the other set-ups.

**Figure 4.**
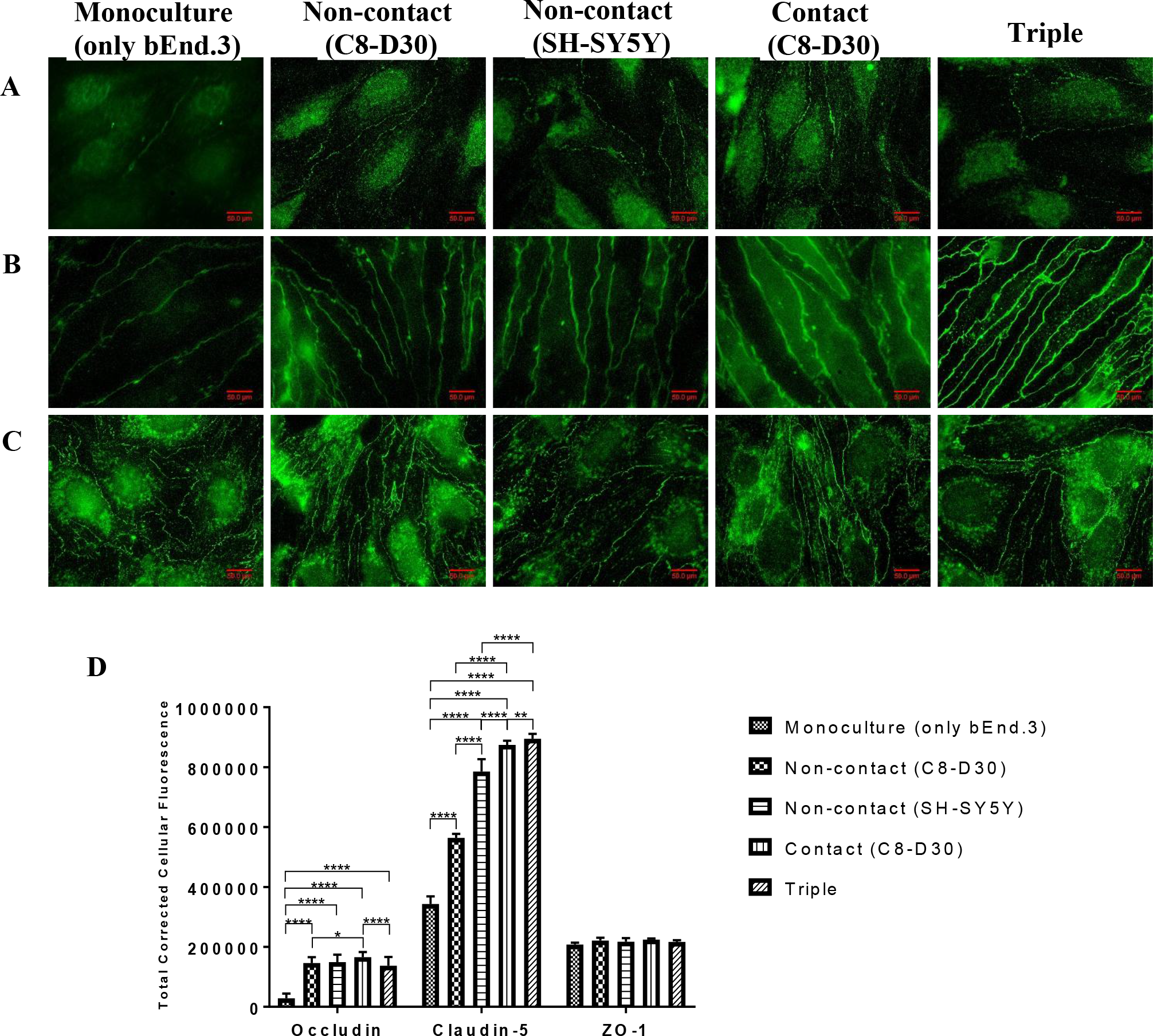
Peripheral localization of Tight Junction proteins of bEnd.3 in different culture models (100X magnification). All TJ proteins A) occludin, B) claudin-5, and C) ZO-1 were detected along the cell contour as continuous lines. The influence of C8-D30 or SH-SY5Y on TJ proteins expression in bEnd.3 were noticeable by the relatively increased fluorescence intensity compared to the monoculture. D) Fluorescence intensity analysis of TJ protein expression in bEnd.3 in different culture models. Total corrected cellular fluorescence (TCCF) were calculated and presented as means ± SD (*n* = 18 – 24). Asterisks indicate significance (* *p*<0.05, ** *p*<0.01, **** *p*<0.0001).

In contrast, localization of claudin-5 was more limited to intercellular junctions outlining clearly the cell borders in bEnd.3 in all culture set-ups. There was no cytoplasmic diffusion of claudin-5 detected. However, in the absence of other cells, bEnd.3 showed irregular, “zipper-like” localization of claudin-5 along the cell periphery. Intensity analysis revealed a significant increase in the peripheral localization of the TJ protein as high as 2.6-fold increase, which was obtained with triple co-culture model compared to the monoculture and significantly high from other model set-ups (Figure 4D). ZO-1 was detected both in the cytoplasm and the cell periphery with a zipper-like appearance in the absence of C8-D30 or SH-SY5Y. However, there was no significant increase in the peripheral distributions of ZO-1. Nonetheless, we observed reduced zipper-like irregularities of the protein, similar to claudin-5, when other cells were introduced in the culture.

### 2.5. Effects of conditioned media on the permeability of bEnd.3 monolayers

Astrocytes and neurons are known to secrete soluble factors which help in the maintenance of BBB integrity (Rubin and Staddon, 1999; Daneman et al., 2009). Hence, the effect of conditioned media derived from astrocytes and neurons on barrier integrity was investigated by culturing bEnd.3 monolayers grown on Transwell system (0.4 μm, polyester, 12-well, Corning Costar) in astrocyte-conditioned media (ACM), neuronal-conditioned media (NCM) or in ACM+NCM in the absence of astrocytes or neurons. Conditioned media were able to induce tightness to bEnd.3 monolayers as shown by the reduced permeability of sucrose and albumin across the monolayers (Figure 5). Papp values of sucrose were significantly decreased in all media conditions with the lowest obtained with ACM+NCM, followed by ACM, and NCM with 139.9-, 20.0-, and 17.2-fold reduction, respectively, compared to the control, but no significant difference was observed between the conditions. ACM and ACM+NCM were also able to decrease the permeability of albumin significantly but not NCM alone with the lowest achieved with ACM+NCM with 16.5-fold reduction compared to the control group.

**Figure 5.**
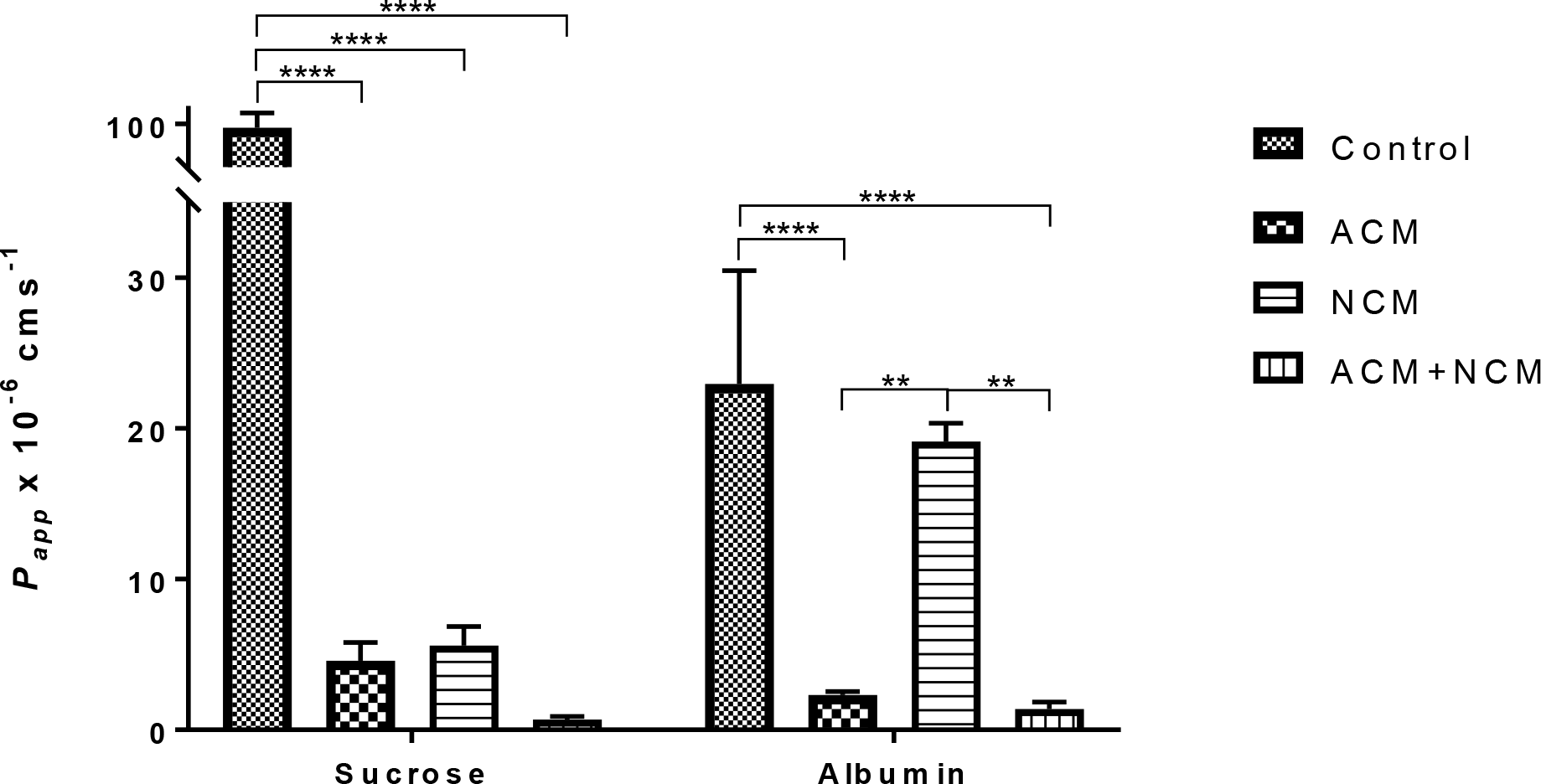
Permeability assessment of bEnd.3 monolayer propagated in conditioned media. bEnd.3 was propagated in complete growth media mixed with astrocyte-conditioned media (ACM), or neuronal-conditioned media (NCM), or ACM+NCM at ratios of 1:1, 1:1, and 2:1:1, respectively. For control condition, bEnd.3 was propagated in complete growth media only (Dulbecco’s Modified Eagle’s medium). Data are presented as means ± SD (*n* = 20 – 24) from 3 independent experiments. Asterisks indicate significance (** *p*<0.01, **** *p*<0.0001).

### 2.6. Effects of conditioned media on TJ proteins expression and peripheral distribution in bEnd.3 cells

We next explored the possible effects of conditioned media on the expression of TJ proteins which may contribute to the increased tightness of bEnd.3 monolayers as seen in the permeability assessment. Western blot analysis revealed that all three TJ proteins expression were not altered as shown in Figure 6. We further investigated the distribution of these proteins in bEnd.3 cells when propagated in conditioned media. The distributions of occludin, claudin-5, and ZO-1 in bEnd.3 cells are shown in Figure 7. Cytoplasmic expression, as well as the junctional distribution of occludin were observed in all conditions. There were significant increase of the peripheral intensities of occludin showing 1.2-, 1.3-, and 1.6-fold increase when propagated in ACM, NCM, and ACM+NCM, respectively, and ACM+NCM condition exerted further significant increase compared to ACM or NCM alone (Figure 7D). Claudin-5 was also detected at the cell periphery, although showing zipper-like irregularities, with minimal cytoplasmic diffusions. However, conditioned media were not able to significantly increase the peripheral localization of claudin-5 as analyzed by the peripheral intensities (Figure 7D). ZO-1 showed similar distribution as occludin with obvious irregularities at the cell junctions. Further intensity analysis of ZO-1 also showed a significant increase with the highest obtained when monolayers were propagated in ACM (2.8-fold increase) and significantly different from the other conditions. In addition, zipper-like irregularities of both claudin-5 and ZO-1 were still observed in monolayers propagated in conditioned media. Although increased peripheral localization was noted, this observation may suggest that conditioned media is not sufficient and the presence of cells physically is important to induce regular and restricted arrangement of TJ proteins at the cell-cell contact.

**Figure 6.**
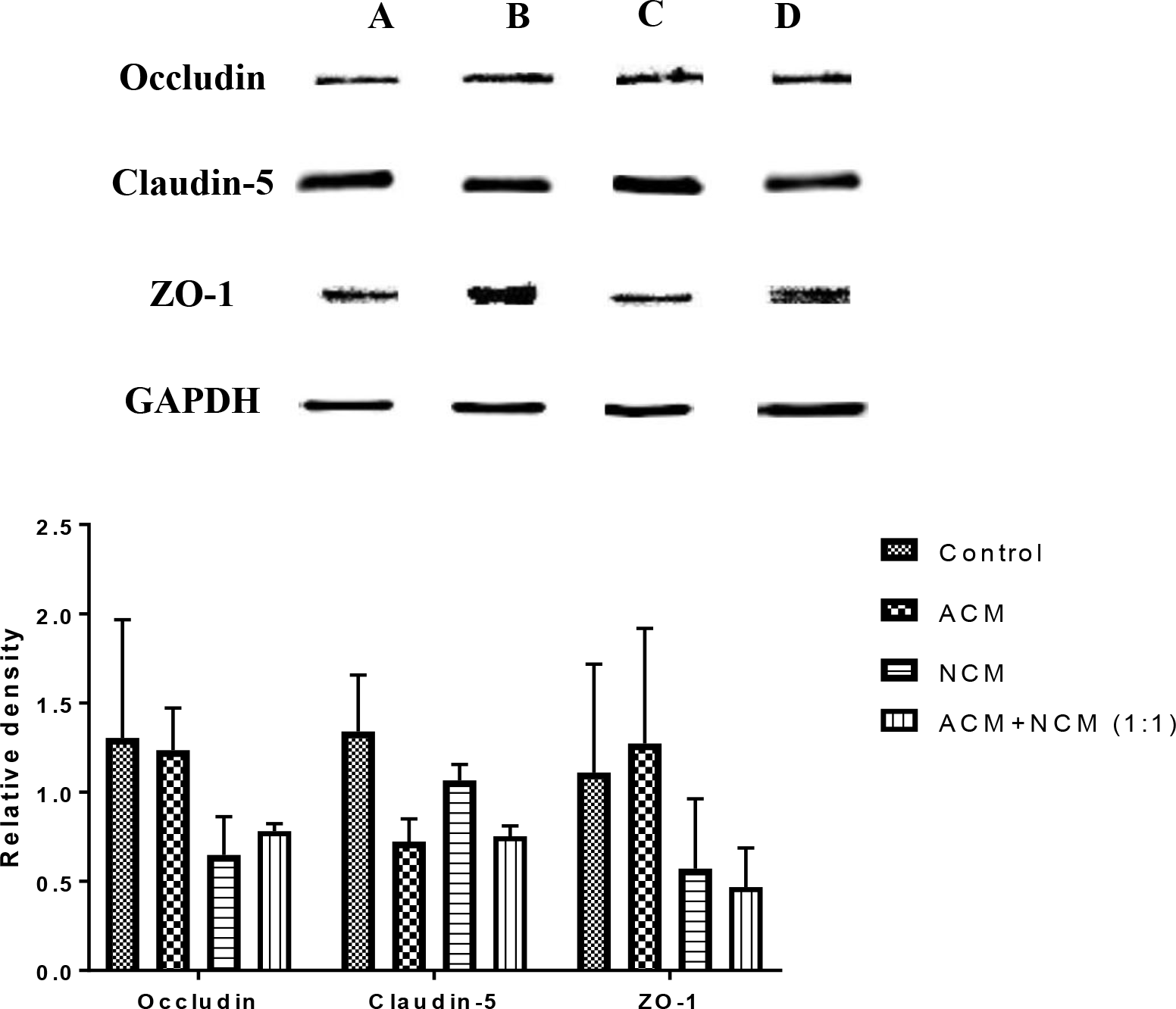
Expression of proteins in bEnd.3 cultured in different conditioned media as determined by Western blotting. A) Control, B) ACM, C) NCM, and D) mixture of ACM and NCM. The relative density with GAPDH as the reference was quantified using ImageJ software. No significant differences in the expression were observed between different conditioned media.

**Figure 7.**
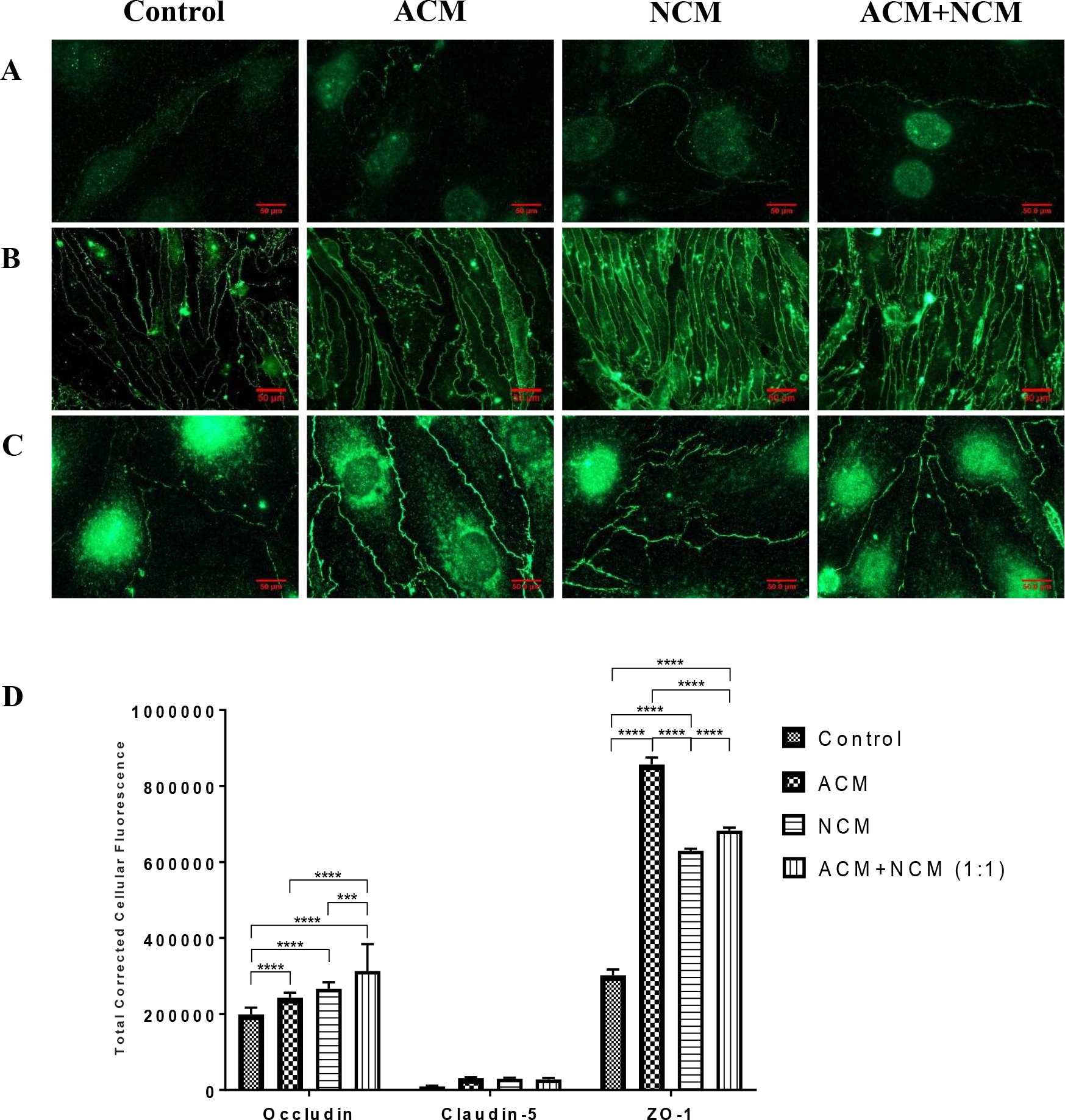
Peripheral localization of Tight Junction proteins of bEnd.3 cultured in different conditioned media (X100 magnification). A) occludin, B) claudin-5, C) ZO-1. bEnd.3 was propagated in complete growth media mixed with astrocyte-conditioned media (ACM), or neuron-conditioned media (NCM), or ACM+NCM at ratios of 1:1, 1:1, and 2:1:1, respectively. For control condition, bEnd.3 was propagated in complete growth media (Dulbecco’s Modified Eagle’s medium). D) Fluorescence intensity analysis of TJ protein expression in bEnd cultured in different conditioned media. Total corrected cellular fluorescence (TCCF) were calculated and presented as means ± SD (*n* = 18 – 24). Asterisks indicate significance (*** *p*<0.001, **** *p*<0.0001).

## 3. Discussion

To date, studies delineating the biochemical and molecular properties of BBB have been performed mainly in cell culture systems employing primary cells originated from various species which include porcine, bovine and rodent (Perrière et al., 2005; Malina et al., 2009; Helms et al., 2010; Nakhlband and Omidi, 2011; Vandenhaute et al., 2011; Cantrill et al., 2012; Patabendige et al., 2013). *In vitro* BBB models using stem cells-derived ECs have also been developed (Lippmann et al., 2012; Boyer-Di Ponio et al., 2014; Cecchelli et al., 2014). These include hPSCs obtained from the inner cell masss of human blastocysts (Thomson et al., 1998) and induced pluripotent stem cells derived from reprogramming somatic cells to a pluripotent state (Takahashi and Yamanaka, 2006; Takahashi et al., 2007). Models developed using hPSC-derived brain cells produced restrictive barrier with peripheral expressions of occludin, claudin-5, and ZO-1, high TEER values reaching 1450 Ωcm^2^ when co-cultured with rat astrocytes (Lippmann et al., 2012), and very low sucrose permeability. Even though primary culture models are considered the best as they can maintain the *in vivo* BBB characteristics at low passage numbers, these models still have the downsides. For example, it has been reported that primary cells are prone to potential contamination by other cell types of the neurovascular unit (NVU) including astrocytes and pericytes which may introduce confounding variables such as different cell monolayer arrangement and involuntary induction of BBB properties to ECs that affect the evaluations of the monolayer integrity developed from a supposed pure EC population. It is also well known that primary cells are difficult due to the slow growth, spontaneous dedifferentiation if the cells are subjected to repeated passages and the limited amount of material such as nucleic acid and proteins that can be produced for molecular and biochemical assays. Considering these potential challenges with primary cell culture models, the current study explored the alternative, easy to maintain BBB model derived from cell lines that may behave closer to the *in vivo* counterparts. Hence, we developed and assessed *in vitro* BBB models using immortalized cell lines employing mouse cell line, bEnd.3 and astrocytes type III, and neuroblastoma of human origin, for its suitability as a BBB model system, and investigated the effect of different cells as well as influence of conditioned media on BBB characteristics and TJ complexes.

The establishment of models required different cell configurations in order to closely mimic the *in vivo* cell arrangements. We developed and tested five different arrangements of cells to best simulate the BBB; monoculture, non-contact co-culture with C8-D30 or SH-SY5Y, contact co-culture with C8-D30 at the basal side of inserts, and triple co-culture, and assessed their tightness using transport study of sucrose and albumin and the regulation of TJ proteins in bEnd.3 in the presence and absence of other cell types. In the permeability assessment, the low permeability of sucrose and albumin confirmed that all culture setups showed a controlled passage of molecules across the barrier, as demonstrated previously (Li et al., 2010; Yuan et al., 2010; Watanabe et al., 2013; Wuest et al., 2013). Our monoculture model showed a high sucrose P_app_ values compared to other bEnd.3 monoculture published in the literature which was reported in the range of 10 to 19 × 10^−6^ cms^−1^ (Omidi et al., 2003; Brown et al., 2007) probably due to the differences in the culture period (6 to 7 days versus 5 days). Nonetheless, our bEnd.3 monoculture was tighter by more than 2-fold compared to few studies that determined the transendothelial permeability of other cell lines. For instance, RBE4 cell monolayers have been reported to produce permeability coefficients for sucrose of 214 × 10^−6^ cms^−1^ (Rist et al., 1997) and permeability value of 195 × 10^−^6 cms^−1^ was obtained with human brain capillary endothelial cell line (BB19) (Kusch-Poddar et al., 2005). Moreover, bEnd.3 monoculture sucrose value (96.1 ± 24.5 × 10^−6^ cms^−1^) was comparable to hCMEC/D3, another immortalized human brain endothelial cell line, which was reported to have 80.7 × 10^−6^ cms^−1^ for sucrose permeability (Al-Shehri et al., 2015).

The addition of astrocytes in a co-culture system was expected to enhance and improve the model since these cells are important for the endothelial cells to acquire a BBB phenotype (Rubin et al., 1991; McAllister et al., 2001; Abbott et al., 2006). Indeed, permeability values of co-culture models with astrocyte (C8-D30) were significantly lower than bEnd.3 monoculture values, which is in agreement with Booth and Kim (2014). Sucrose permeability values of co-culture models with astrocytes obtained in this study; 1.1 ± 0.2 and 1.8 ± 0.6 × 10^−6^ cms^−1^ for non-contact and contact, respectively, were comparable to that of the primary mouse or rat brain endothelial cells which reported permeability values of sodium fluorescein of 3.5 to 6.5 × 10^−6^ cms^−1^ (Shayan et al., 2007; Wuest et al., 2013) and 0.5 to 8.2 × 10^−6^ cms^−1^ (Weidenfeller et al., 2007; Sumi et al., 2010; Mallareddy et al., 2012), respectively. Even though the study used sodium fluorescein as the paracellular marker instead of sucrose, it is expected that the values will not deviate greatly as there is a little difference in the molecular weight of sucrose (342.3Da) and sodium fluorescein (376Da), allowing comparisons to be made. Albumin values could not be compared directly with other literature since studies employing bEnd.3 cells as their model did not assess the permeability of albumin. However, it was comparable, although our co-culture values were slightly high, to primary rat brain endothelial cells (Hiu et al., 2008) and hCMEC/D3 (Hsuchou et al., 2010).

Co-culturing endothelial cells with astrocytes are known to increase monolayer resistance compared to endothelial cell grown in monoculture (Gaillard et al., 2001; Jeliazkova-Mecheva and Bobilya, 2003; Garcia et al., 2004), which is in agreement with our findings. This supports earlier observations by Goldstein (1988) who described that the presence of astrocytes in endothelial cells co-cultures increases the complexity and development of TJs as well as the expression of specific BBB markers. On the contrary, our findings showed the opposite to that observed in several studies (Omidi et al., 2003; Weksler et al., 2005; Cucullo et al., 2008; Wuest et al., 2013), reporting no additional benefit of co-culturing with astrocytes which may have been due to the type of astrocyte employed and/or serum supplementation during propagation. It is noteworthy that variation in the types of astrocytes used in co-culture systems may contribute to the discrepancies. C8-D30 used in this study was reported to resemble a phagocytic or reactive astrocyte in tissue and is not commonly associated with the brain blood vessels *in vivo* unlike astrocyte type I (An et al., 2011). Nonetheless, we have shown this type of astrocyte was able to induce BBB properties on bEnd.3 which may reflect similar ability to astrocyte type I. The findings by Hatherell et al. (2011) support our observation in which two different types of astrocytes derived from either cerebral or cerebellar exerted the same influence on the TJ resistance and no significant differences were observed suggesting that regardless of the origins, astrocytes play a role in the induction of BBB properties.

In the presence of neurons as in the non-contact model, we observed a significant decrease in the Papp of sucrose but not albumin across the model which suggest that neurons play a role in the induction of BBB characteristics. Neurons have been shown to decrease sucrose leakage across *in vitro* BBB (Schiera et al., 2005), possibly through the regulation of occludin localization (Savettieri et al., 2000; Schiera et al., 2003). Neurons are also reported to have the ability to induce BBB properties in brain endothelial cells (Tontsch and Bauer, 1991; Cestelli et al., 2001) in which Cestelli et al. (2001) reported that RBE4 co-cultured with rat primary neurons reduced the transmonolayer flux of dopamine. Furthermore, SH-SY5Y has been reported to be able to induce BBB properties in PBEC and RBE4 (Toimela et al., 2004; Freese et al., 2014).

Several triple co-culture BBB models were reported previously which employed combinations of primary brain endothelial cells or cell lines, astrocytes, neurons, and pericytes (Xue at al., 2013; Adriani et al., 2015; Thomsen et al., 2015; Vandenhaute et al., 2011; Takata et al., 2013). However, no permeability coefficient values can be found determined on bEnd.3 in triple co-culture to compare with the results obtained in the present study. Likewise, there are no sucrose permeability values found to compare with and again we compare our values with that of the sodium fluorescein. Several studies indicated a low sodium fluorescein permeability across their BBB models developed using primary rat brain endothelial cells co-cultured with pericytes and astrocytes with values in the range between 0.58 and 6.4 × 10^−6^ cms^−1^ (Imamura et al., 2011; Wilhelm et al., 2011; Hellinger et al., 2012; Takata et al., 2013; Jahne et al., 2014). Triple co-culture consisting of RBE4, pericytes, and astrocytes also showed a low permeability to sodium fluorescein of 3.9 ± 0.2 × 10^−6^ cm^−1^ (Nakagawa et al., 2009). We demonstrated that co-culturing of bEnd.3 with C8-D30 and SH-SY5Y decreased the flux of sucrose significantly to 0.56 ± 0.16 × 10^−6^ cms^−1^. In contrast, albumin permeability value of our model was higher than these established models which were reported as low as 0.18 × 10^−6^ cms^−1^ (Takata et al., 2013), while ours was 3.4 ± 0.2 × 10^−6^ cms^−1^. These differences in values may be attributed by the species difference (rat vs mouse), pore sizes of the inserts used and the use of coating agents, typically collagen and fibronectin, on inserts during seeding.

Nevertheless, the permeability of our triple co-culture to sucrose does compare well with data in the literature which used primary bovine and porcine endothelial cell monolayers. These cells are considered to be the best cell models for BBB development due to their highly restrictive nature. For instance, primary bovine brain microvascular endothelial cells consistently produced values in the range of 5 and 23 × 10^−6^ cms^−1^ (Johnson and Anderson, 1999; Descamps et al., 2003; Lockman et al., 2003; Nakhlband and Omidi, 2011), while values ranging between 0.34 and 12.1 × 10^−6^ cms^−1^ have been reported for porcine brain endothelial cells (Lohmann et al., 2002; Kusch-Poddar et al., 2005; Smith et al., 2007; Patabendige et al., 2013. Hence, our data may suggest that our triple co-culture model is as restrictive and comparable to BBB models constructed using primary cells. We also could not dismiss the fact that inserts may also restrict the passage of molecules in the absence of cells. In the contact and triple co-culture, endothelial cells and C8-D30 were separated by a polyester membrane with a thickness of 10 μm and pore sizes or 0.4 μm and thus, it is likely that the enhancement of the barrier properties was a result from a direct contact between astrocytic processes and bEnd.3. In fact, it has been demonstrated that pore sizes of approximately 0.4 μm in diameter allowed astrocyte processes to go through the pores and interact with endothelial cells on the apical side while restricting the movement of astrocytes to the seeded endothelial side (Ma et al., 2005).

Astrocytes to neurons ratio may play a role in contributing to the restrictiveness of BBB and phenotype induction. We used an equal ratio of astrocytes and neurons in the development of triple co-culture model as the optimal ratio of these cells necessary to induce BBB properties *in vitro* was not widely described in literature. However, several *in vivo* studies have reported that in the cerebral cortex from which endothelial cells were obtained, glial cells always outnumber neurons. For examples, in the adult rat cortex, glial cells were found to be more abundant than neuron in a 1.5:1 ratio (Herculano-Houzel and Lent, 2005), while the human cortex has a glial to neuron ration of 3.76:1 (Azevedo et al., 2009). Our study did not investigate the optimum ratio between astrocytes and neurons population but used a astrocytes: neurons ratio of 1 : 1 in the triple co-culture model. We showed that this ratio was able to decrease the permeability of sucrose by 172.2-fold, which may suggest that the abundance of particular cells does not play many roles in the tightness of BBB model at least *in vitro*.

We further investigated the roles of TJ proteins which may contribute to the tightness of these five BBB models. Occludin was observed to be localized in the cytoplasm of the cells in the monoculture, similar observations by Song and Pachter (2003) and Brown et al. (2007). We also detected occludin in the cell periphery in some cells lining the cell-cell junctions. While both claudin-5 and ZO-1 were strongly expressed in the vicinity of cell borders, which is in contrast to the findings by Yang et al. (2017) who observed weak and diffused claudin-5 in their study. This discrepancy may be due to the shorter incubation period and low cell seeding concentration compared to our study (3 to 4 days vs 5 days incubation and 3 × 10^4^ vs 5 × 10^4^ cells/well). Co-culturing of bEnd.3 with C8-D30 or SH-SY5Y increased the peripheral localization of occludin and claudin-5, although the protein expressions as determined by Western blot remained unaltered. bEnd.3/ astrocyte co-culture was reported to have no effect on the expression of TJ proteins (Li et al., 2010). Neurons were reported to induce peripheral localization of occludin in endothelial cells after several days of co-culturing (Savettieri *et al.* 2000). Non-contact co-culture of brain endothelial cells with SH-SY5Y significantly increased the tightness of both endothelial cells and strong expressions of occludin, claudin-5, and ZO-1 in PBEC (Freese et al., 2014). bEnd.3 in the triple co-culture model, which exhibits the tightest paracellular and transcellular barrier for sucrose and albumin among the five models tested in our experiments, expressed a relatively high level of TJ proteins occludin, claudin-5, and ZO-1. The claudin-5 and ZO-1 localization were also more restricted to the endothelial junctions in the triple co-culture than in the monoculture. The elevated claudin-5 expression and junctional localization are particularly significant in this model since this TJ protein is the only one which has been associated directly to the restrictiveness of BBB *in vivo* (Nitta et al., 2003, Liu et al., 2015).

In addition, there is a possibility that the increase in the BBB restrictiveness might be due to the astrocyte- or neuronal-mediated discharge of short-lived factors that are capable of passing the Transwell inserts in order to regulate the BBB properties. Our study found ACM and NCM to be equally effective at improving the barrier properties of bEnd.3 monolayers. Propagating bEnd.3 with either ACM, NCM or in a mixture significantly reduced the permeability of sucrose and albumin as high as 139.9 and 16.5-fold, respectively, obtained with mixtures of ACM and NCM. However, no significant reduction of albumin flux was observed during propagation in the presence of NCM. Studies by Rist et al. (1997) and Lagrange et al. (1999) reported that exposure of rat brain endothelial cell line, RBE4, to astroglial factors decreased the sucrose permeability across the monolayer to a range between 38 and 68 × 10^−6^ cms^−1^. Furthermore, the addition of conditioned media especially that was derived from astrocytes was demonstrated to reduce the permeability of sucrose and FITC-conjugated albumin (Rubin et al., 1999) as well as propidium iodide (Kuo and Lu, 2011) across bovine and human BBB models. Increased peripheral localization of occludin and ZO-1, but not claudin-5, in bEnd.3 was also observed when the endothelial cells were propagated in conditioned media, although no significant alterations in these TJ proteins expression were detected. This effect has been described by Siddharthan et al. (2007) in which conditioned media particularly ACM was able to stimulate the expression of ZO-1.

## 5. Conclusion

We successfully developed 5 different models with different configurations using bEnd.3, C8-D30 and SH-SY5Y as brain endothelial cells, astrocyte and neuron cell models, respectively, which we believe to be the first study to combine these cells in the establishment of *in vitro* BBB models. We observed that contact and triple co-cultures provide reliable and reproducible *in vitro* BBB models possessing the properties very close to those demonstrated *in vivo*. Triple co-culture model showed the lowest permeability to sucrose and albumin indicating a restrictive barrier which is comparable to other established triple co-culture employing primary rat brain endothelial cells, pericytes and astrocytes. We also observed an increase in the TJ protein expression particularly occludin in bEnd.3 in the presence of C8-D30 or SH-SH5Y, and co-culturing lead to the increase of peripheral localization of occludin and claudin-5. We also found that conditioned media especially ACM could induce BBB properties as shown by the low permeability of sucrose and albumin across bEnd.3 monolayers propagated in these conditioned media by regulating the localization of occludin and ZO-1 to the cell-cell junctions, although no significant increase in the TJ proteins expression was observed. This indicates that astrocyte and neuronal factors play a role in the induction of BBB phenotype.

## 4. Experimental procedures

### 4.1. Cell culture

Mouse brain endothelial cells (bEnd.3) CRL2299, Astrocyte type III (C8-D30) CRL2534, and Neuroblastoma SH-SY5Y CRL2266 were purchased from American Type Culture Collection (ATCC). All cells were grown in Dulbecco’s modified Eagle’s medium (DMEM) (Sigma) containing 4500 mg glucose L^−1^, 110 mg sodium pyruvate L^−1^ and sodium bicarbonate, supplemented with 10% fetal calf serum, 2 mM L-glutamine and 100 I.U. penicillin-streptomycin ml^−1^. One-half of the medium was changed every third or fourth day. Cells were maintained under constant conditions of 37°C, 5% CO_2_ and a humidified atmosphere in a cell culture incubator.

For the study on the effect of conditioned media, NCM and ACM were collected when passaging SH-SY5Y and C8-D30, respectively, stored in −20°C until further use. Briefly, 5.0 × 10^4^ bEnd.3 cells were plated on the apical side of Transwell inserts as a monolayer in either DMEM:NCM or DMEM:ACM media at a ratio of 1:1. The effect on the bEnd.3 integrity of a third mixture composing of growth medium (DMEM), ACM, and NCM at a ratio of 2:1:1 was also examined and assessed.

### 4.2. Construction of *in vitro* BBB models

#### 4.2.1. Monoculture

bEnd.3 cells (5.0 × 10^4^ cells) were plated on the apical of Transwell inserts (0.4 μm, polyester, Corning Costar). Cells were grown to confluency up to 5 days.

#### 4.2.2. Non-contact co-culture

This system was established following Molino et al. (2014) protocol with modifications. bEnd.3 cells (5.0 × 10^4^ cells) were seeded on the apical side of the Transwell inserts and C8-D30 or SH-SY5Y (1.0 × 10^5^ cells) were seeded at the bottom of a 12-well plate.

#### 4.2.3. Contact co-culture

The system was established according to Zhang et al. (2011) with modifications. However, this model was established with astrocytes only as neurons are not in direct contact with endothelial cells *in vivo*. Briefly, C8-D30 were seeded directly (5.0 × 10^4^ cells) on the basal side of the Transwell inserts and allowed to attach and proliferate on the insert for 2 days before being inverted. bEnd.3 (5.0 × 10^4^ cells) were then seeded at seeding concentration of on the apical side. The culture was further incubated up to 5 days.

#### 4.2.4. Triple co-culture system

Triple culture system was established following Takata et al. (2013) protocol with modifications. The system combined three cells in one culture system. In this system, SH-SY5Y (5.0 × 10^4^ cells) was propagated at the bottom of a 12-well plate. Inserts with confluent C8-D30 grown on the basal side were transferred into the well containing SH-SY5Y. bEnd.3 (5.0 × 10^4^ cells) was then seeded on the apical side and cells were maintained for a further 5 days.

### 4.3. *In vitro* evaluation of the barrier function

The barrier function of the developed models was evaluated using transport study determining the permeability of sucrose and albumin as paracellular and transcellular markers, respectively. Transport studies were performed on each culture set up according to Zhang et al. (2011) method with modifications. Briefly, inserts containing endothelial monolayer were washed in excess PBS before transferring the inserts in wells containing serum-free, phenol red-free DMEM in the basal compartment. 0.5 ml of compounds (50 mgml^−1^) dissolved in serum-free, phenol red-free DMEM were added in the apical compartment and were incubated for 15, 30, 60, 90, 120 and 150 minutes at 37°C, 5% CO_2_ and a humidified atmosphere in a cell culture incubator. 1 ml of media was extracted from the basal compartment at the end of each time point and measured using UV/VIS spectrometry (PerkinElmer Lambda 25) to determine the concentration of transported compounds. The absorbance of sucrose was measured at 210 nm and 270 nm, while the absorbance of albumin was measured at 279 nm and 290.5 nm. The compound concentration of the extracted samples was calculated based on the corresponding calibration curves. Subsequently, P_app_ values indicating the rates at which compounds diffuse to the basal compartment of the Transwell were calculated using the following equation according to Hubatsch et al. (2007):

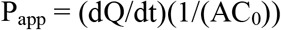

Where dQ/dt is the steady-state flux (mgml^−1^s^−1^), A is the surface area of the insert membrane (1.12 cm^2^), and C_0_ is the initial concentration in the donor chamber in mgml^−1^.

### 4.4. Western Blot analysis

Cellular extracts were prepared in RIPA buffer containing 50 mM Tris, 150 mM NaCl, 1% NP40, 1 mM EDTA, 0.25% sodium deoxycholate, and 1% protease inhibitor cocktail (SIGMA).

The total protein concentration in the cell lysates was determined using a BCA Protein assay (Pierce, Rockford, IL, USA). Equivalent amounts of protein from each sample were subjected to SDS-PAGE using 10% gel followed by electrotransfer to a nitrocellulose membrane. The nitrocellulose membranes were blocked in TBS buffer 0.1% Tween 20 containing 5% nonfat dry milk for 1 hour. Occludin, claudin-5, ZO-1, and GAPDH were detected using antibodies against occludin (1:2000, Abcam), claudin-5 (1:2000, Abcam), ZO-1 (1:500, Merck), and GAPDH (1:2000, Abcam) with overnight incubation at 4°C, followed by incubation with HRP-conjugated anti-rabbit IgG antibodies. Clarity™ Western ECL substrate was used for protein detection and imaging was performed using Molecular Imager VersaDoc™ MP 4000 System (Bio-rad). The relative intensity of the individual proteins was expressed as the ratio of GAPDH and the corresponding total protein.

### 4.5. Immunofluorescence assay

bEnd.3 on Transwell inserts were washed with PBS 3 times to remove medium before fixing in ice-cold acetone for 1 minute. Inserts were then washed with PBS twice to remove excess acetone. 200μl of permeabilization buffer (0.2% Triton-X100, 2% BSA) was applied to each insert and incubated for at least 30 minutes. Third washing was done and primary antibodies against occludin (1:1000, Abcam), claudin-5 (1:1000, Abcam), and ZO1 (1:500,Merck) diluted in antibody signal enhancer (10mM glycine, 0.05% Tween20, 0.1% Triton X-100, 0.1% hydrogen peroxide in PBS) (Rosas-Arellano et al., 2016) was applied onto each sample and were incubated for 1 hour at room temperature. This was followed by incubating inserts with a conjugated secondary antibody (1:1000 dilution in permeabilization buffer) for 1 hour at room temperature. Subsequently, inserts were washed with distilled water three times and mounted in glycerol/PBS (1:1). Imaging was done using epifluorescence microscope Nikon Eclipse 90i equipped with a Nikon DS-Fi1 camera.

To quantify proteins expression based on the fluorescence intensity, a single in-focus plane was acquired according to McCloy et al. (2014). An outline was drawn around expressed protein in each cell and analyzed using Fiji/ImageJ software to obtain the area and mean fluorescence measurements, as well as several adjacent background readings. The total corrected cellular fluorescence (TCCF) was calculated according to the formula below:

TCCF = integrated density – (area of selected cell x mean fluorescence of background readings)

### 4.6. Statistical analysis

All results were expressed as means ± SD from three or more independent experiments. Statistical significance was assessed by 2-way ANOVA multiple comparison tests. A *p* value < 0.05 was considered as significant. All statistical analyses were performed using GraphPad Prism 7.0 for Windows (GraphPad Software, San Diego, California, USA.

## Funding

This research did not receive any specific grant from funding agencies in the public, commercial, or not-for-profit sectors.

## Acknowledgements

This work is supported by Universiti Brunei Darussalam. Idris F is a recipient of the Graduate Research Scholarships (GRS), Universiti Brunei Darussalam.

